# **McImpute**: Matrix completion based imputation for single cell RNA-seq data

**DOI:** 10.1101/361980

**Authors:** Aanchal Mongia, Debarka Sengupta, Angshul Majumdar

## Abstract

**Motivation:** Single cell RNA sequencing has been proved to be revolutionary for its potential of zooming into complex biological systems. Genome wide expression analysis at single cell resolution, provides a window into dynamics of cellular phenotypes. This facilitates characterization of transcriptional heterogeneity in normal and diseased tissues under various conditions. It also sheds light on development or emergence of specific cell populations and phenotypes. However, owing to the paucity of input RNA, a typical single cell RNA sequencing data features a high number of dropout events where transcripts fail to get amplified.

**Results:** We introduce mcImpute, a low-rank matrix completion based technique to impute dropouts in single cell expression data. On a number of real datasets, application of mcImpute yields significant improvements in separation of true zeros from dropouts, cell-clustering, differential expression analysis, cell type separability, performance of dimensionality reduction techniques for cell visualization and gene distribution.

**Availability and Implementation:** https://github.com/aanchalMongia/McImpute_scRNAseq

## Introduction

In contrast to traditional bulk population based expression studies, single cell transcriptomics provides more precise insights into functioning of individual cells. Over the past few years this powerful tool has brought in transformative changes in the conduct of functional biology [39]. With single cell RNA sequencing (scRNA-seq) we are now able to discover subtypes within seemingly similar cells. This is particularly advantageous for characterizing cancer heterogeneity [27, 34], identification of new rare cell type and understanding the dynamics of transcriptional changes during development [1, 33, 40].

Despite all the goodness, scRNA-seq technologies suffer from a number of sources of technical noise. Most important of these is insufficient input RNA. Due to small quantities transcripts are frequently missed during the reverse transcription step. As a direct consequence, these transcripts are not detected during the sequencing step [12]. Often times the lowly expressed genes are the worst hit. Excluding these genes from analysis may not be the best solution as many of the transcription factors and cell surface markers are sacrificed in this process [38]. Added to that, variability in dropout rate across individual cells or cell types, works as a confounding factor for a number of downstream analyses [18, 30]. Hicks and colleagues [8] showed, on a number of scRNA-seq datasets, that the first principal components highly correlate with proportion of dropouts across individual transcriptomes. In summary, there is a standing need for efficient methods to impute scRNA-seq datasets.

Very recently, efforts have been made to devise imputation techniques for scRNA-seq data. Most notable of among these are MAGIC [38], scImpute [19] and drImpute [15]. MAGIC uses a neighborhood based heuristic to infer the missing values based on the idea of heat diffusion, altering all gene expression levels including the ones not affected by dropouts. On the other hand, scImpute first estimates which values are affected by dropouts based on Gamma-Normal mixture model and then fills the dropout values in a cell by borrowing information of the same gene in other similar cells, which are selected based on the genes unlikely affected by dropout events. Overall performance of scImpute has been shown to to be superior to MAGIC. Parametric modeling of single cell expression is challenging due to our lack of knowledge about possible sources of technical noise and biases [30]. Moreover, there is clear lack of consensus about the choice of probability density function. Another method, Drimpute, repeatedly identifies similar cells based on clustering, and performs imputation multiple times by averaging the expression values from similar cells, followed by averaging multiple estimations for final imputation. We propose mcImpute, an imputation algorithm for scRNA-seq data which models gene expression as a low rank matrix and sprouts in values in place of dropouts in the process of recovering the full gene expression data from sparse single cell data. This is done by applying soft-thresholding iteratively on singular values of scRNA-seq data. One of the salient features of mcImpute is that it does not assume any distribution for gene expression.

We first evaluate the performance of mcImpute in separating “true zero” counts from dropouts on single cell data of myoblasts [35] (We call it Trapnell dataset). On the same dataset, we assess the impact of imputation on differential genes prediction. We further investigate mcImpute’s ability to recover artificially planted missing values in a single cell expression matrix of mouse neurons [37].Accurate imputation should enhance cell type identity i.e., transcriptomic similarity between cells of identical type. We therefore quantify cell type separability as a metric and assess its improvement. In addition to these, we also test the impact of imputation on cell clustering. Four independent real datasets Zeisel, Jurkat-293T, Preimplantation and Usoskin ( [43], [41], [40], [37]), for which cell type annotations are available and one dataset, Trapnell, ( [35]) for which bulk RNA-seq data has been provided (required for validation of differential genes prediction and separation of “true zeros” from dropouts), are used for this purpose. mcImpute clearly serves as a crucial tool in scRNA-seq pipeline by significantly improving all the above mentioned metrics and outperforming the state-of-the-art imputation methods in majority of experimental conditions.

With the advent of droplet based, high-throughput technologies [23, 41], library depth is being compromised to curb the sequencing cost. As a result, scRNA-seq datasets are being produced with extremely high number of dropouts. We believe that great performance, will make mcImpute the method of choice for imputing scRNA-seq data.

## Results/Discussion

We performed numerous experiments to evaluate the efficacy of our proposed imputation technique comparing mcImpute with a number of existing imputation methods for single cell RNA data: scImpute, drImpute and MAGIC.

### Dropouts vs true zeros

The inflated number of zero counts in scRNA-seq data could either be biologically driven or due to lack of measurement sensitivity in sequencing. The transcript which is not detected because of failing to get amplified in sequencing step, essentially corresponds to a “false zero” in the finally observed count data and needs to be imputed. A reasonable imputation strategy which has this discriminating property should keep the “true zero” counts (where the genes are truly expressed and have no transcripts from the beginning) untouched, while at the same time attempt to recover the dropouts.

We investigate the performance of mcImpute in distinguishing “true zero” counts from dropouts on Trapnell data [35], for which the bulk-counterpart was available and hence, we could pull out low-to-medium expression genes from the corresponding bulk data for validation. The fraction of zero counts were observed for genes with expression ranging from zero to 500 for unimputed and imputed gene-expression data. It should be noted that an imputed count value ranging from 0-0.5 is taken as an imputed zero, rendering minor flexibility to all imputation techniques.

Given the nature of this analysis, gene filtering in single cell expressions has been skipped. DrImpute could not be taken into account since we could not programatically mute the gene filtering step in its pipeline.

**Fig 1.**
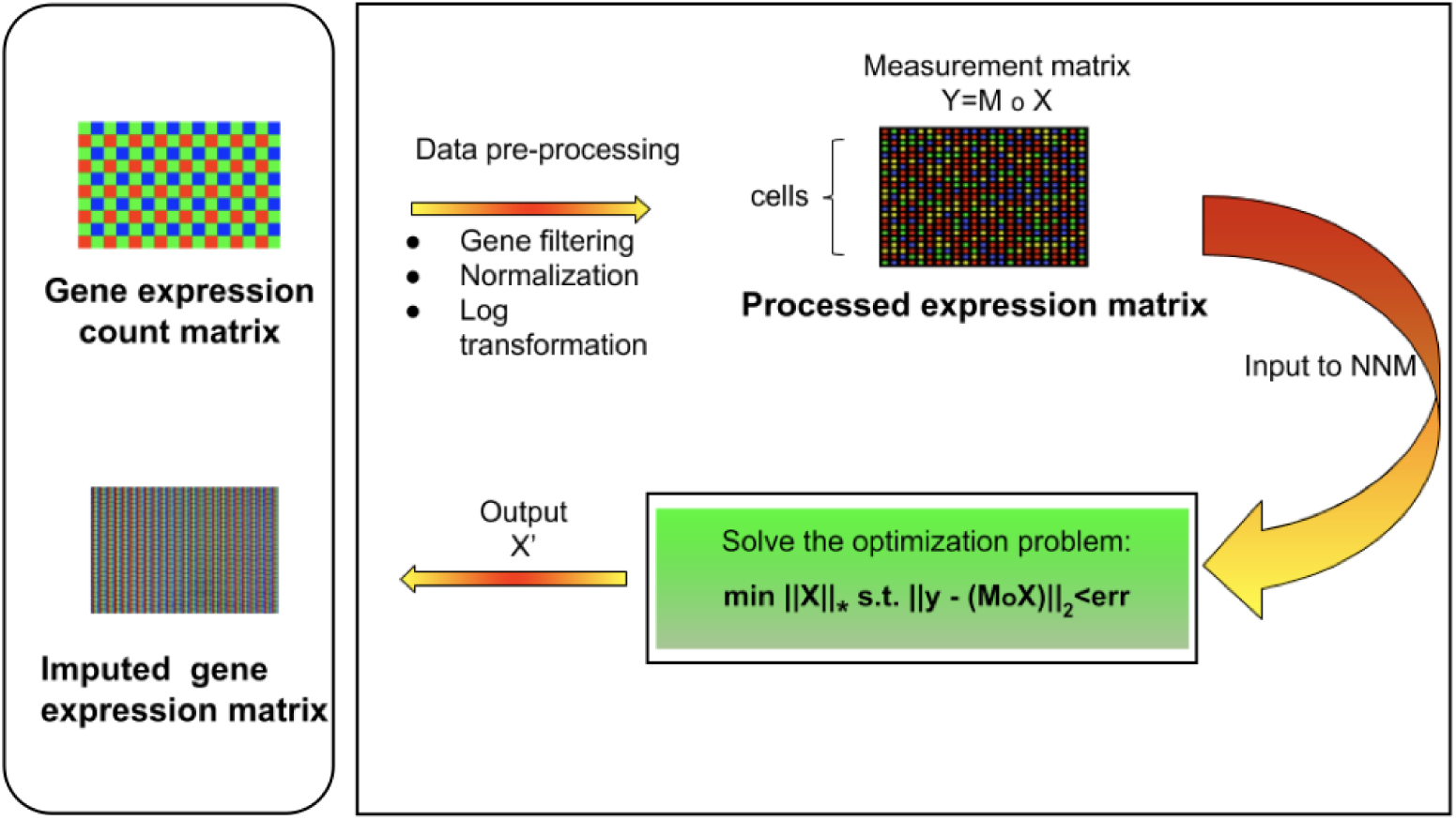
**Overview of mcImpute framework** for imputing single cell RNA sequencing data. Raw read counts were filtered for significantly expressed genes and then normalized by Library size. Then, the expression data was Log2 transformed (after adding a pseudo count of 1). This pre-processed expression matrix (Y) is treated as the measurement/observation matrix (and fed as input to mcImpute) from which the gene expressions of the complete matrix (X) need to be recovered by solving the non-convex optimization problem minimizing nuclear norm of expression matrix.

We observe (figure 2.(e)) that with low expression genes, all imputation strategies successfully impute the “true zeros” while, as the gene expression amplifies, un-imputed matrix still exhibits large fraction of zeros, which essentially correspond to dropouts and only mcImpute and scImpute are able to curtail the fraction of zeros, thus recovering the dropouts back.

**Fig 2.**
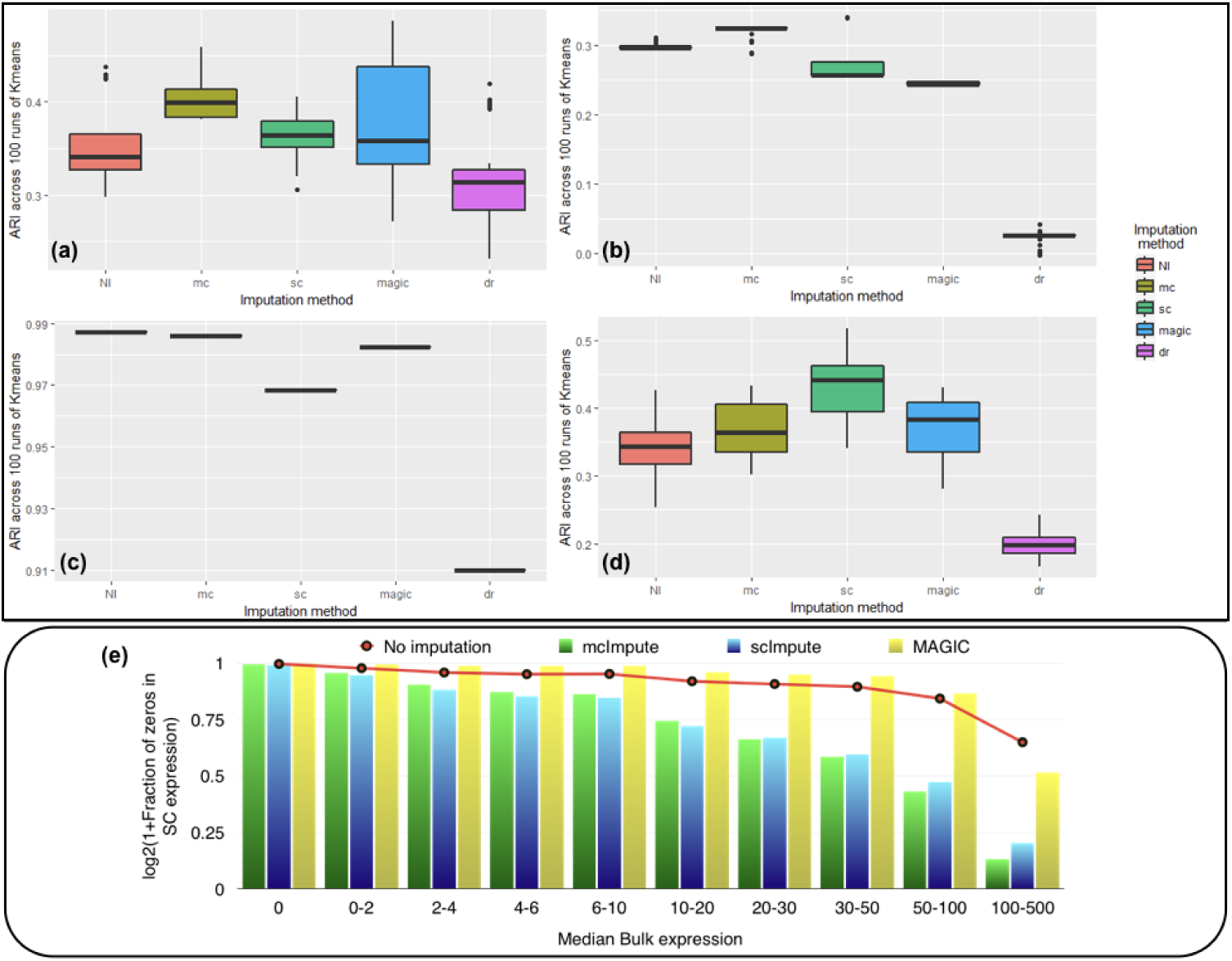
McImpute shows remarkable improvement in clustering of single cells and separation of “true zeros” from dropouts **(a)-(d) Boxplots showing the distribution of ARI** calculated on 100 runs of k-means clustering algorithm on first two principal components of single cell expression matrix for datasets **(a)** Zeisel **(b)** Usoskin **(c)** Jurkat-293T and **(d)** Preimplantation. **(e) Separation of “true zeros” from dropouts**: Plot showing fraction of zero counts (values between 0 and 0.5) in single cell expression matrix against the median bulk expression. The genes are divided into 10 bins based on median bulk genes expression (first bin corresponds to zero expression genes)

### Matrix recovery

In this set of experiments, we study the choice of matrix completion algorithm – matrix factorization (MF) or nuclear norm minimization (NNM). Both the algorithms have been explained in section **Materials and Methods**.

The experiments are carried out on the processed Usoskin dataset [37]. We artificially removed some counts at random (sub-sampling) in the data to mimic dropout cases and used our algorithms (MF and NNM) to impute the missing values. Figure 3.(a)-(c) and **table S2** show the variation of Normalized Mean Squared Error (NMSE), Root Mean Squared Error (RMSE) and Mean Absolute Error (MAE) to compare our two methods for different sub-sampling ratios. This is the standard procedure to compare matrix completion algorithms [11, 25].

**Fig 3.**
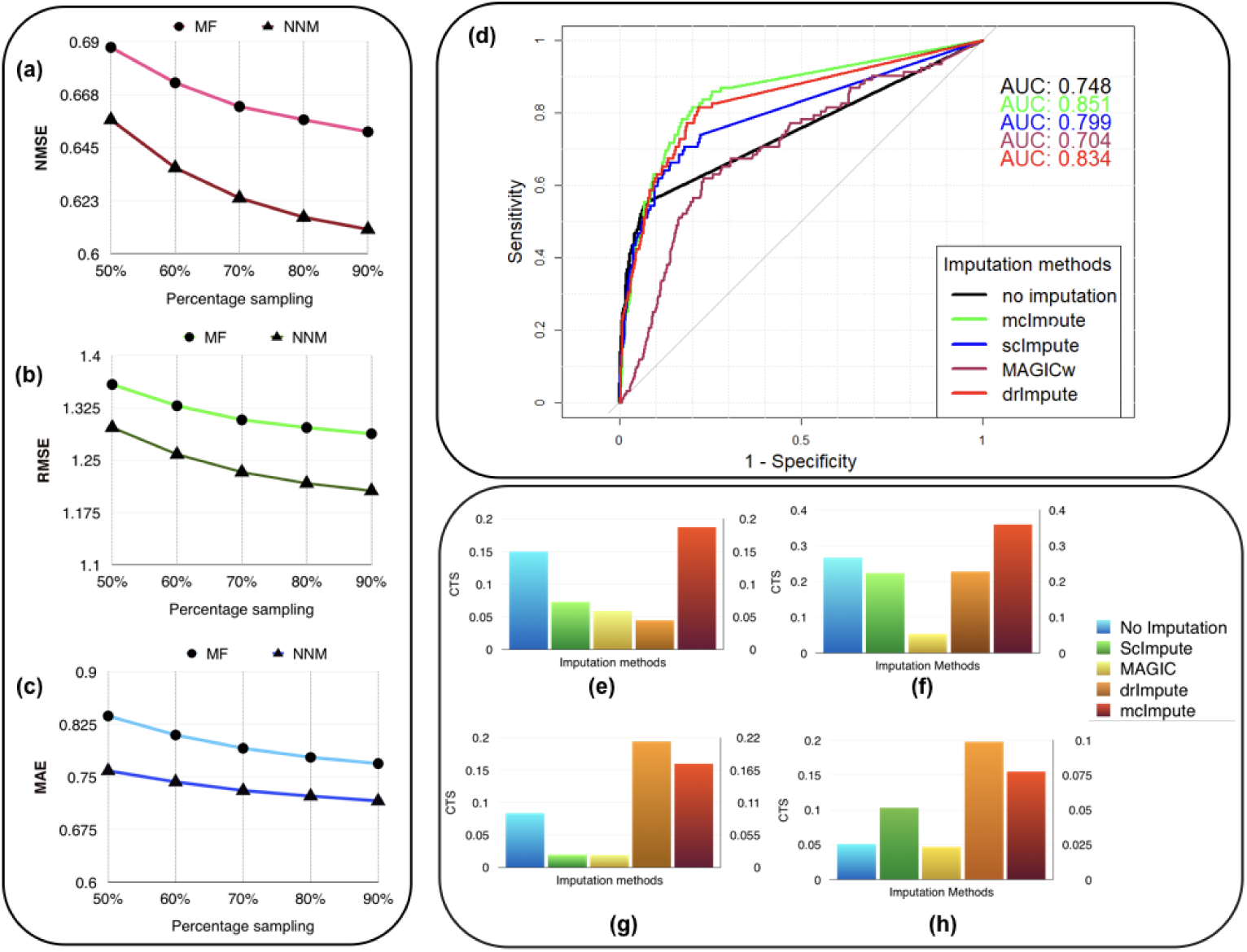
McImpute recovers the original data from their masked version with low error, performs best in prediction of differentially expressed genes and significantly improves CTS score. Variation of **(a)** NMSE, **(b)** RMSE and **(c)** MAE with sampling ration using ARM and MF on Usoskin dataset showing NnM performing better than MF algorithm. **(d)** ROC curve showing the agreement between DE genes predicted from scRNA and matching bulk RNA-Seq data [35]. DE calls were made on expression matrix imputed using edgeR. **(e)-(h)** 2D-Axis bar plot depicting improvement in Cell type separabilities between (e) Jurkat and 293T cells from Jurkat-293T dataset; (f) 8cell and BXC cell types from Preimplantation dataset; (g) S1pyramidal and Ependymal from Zeisel dataset and (h) NP and NF cells from Usoskin dataset. Refer **table S3** for absolute values.

We are showing the results for Usoskin dataset, but we have carried out the same analysis for other datasets and the conclusion remained the same. We find that the nuclear norm minimization (NNM) method performs slightly better than the matrix factorization (MF) technique; so we have used NNM as the workhorse algorithm behind mcImpute.

### Improvement in clustering accuracy

Correct interpretation of single cell expression data is contingent on accurate delineation of cell types. Bewildering level of dropouts in scRNA-seq data often introduces batch effect, which inevitably traps the clustering algorithm. A reasonable imputation strategy should fix these issues to a great extent. In a controlled setting we therefore examined if the proposed method enhanced clustering outcomes. For this, we ran *K*-means on first 2 principal component genes of log transformed expression profiles featured in each dataset. Since the prediction from this clustering algorithm tends to change with the choice of initial centroids, which are chosen at random, we analyze the results on 100 runs of k-means to get reliable and robust results. We set the number of annotated cell types as the value of *K* for every data. Adjusted Rand Index (ARI) was used to measure the correspondence between the clusters and the prior annotations.

McImpute based re-estimation best separates the four groups of mouse neural single cells from Usoskin dataset and brain cells from Zeisel dataset, and clearly shows comparable improvement on other datasets too (figure 2.(a)-(d), **table S3**). Striking difference between Jurkat and 293T cells made them trivially separable through clustering, leading to same ARI across all 100 runs. Still, mcImpute was able to better maintain the ARI in comparison to other imputation methods.

### Improved differential Genes prediction

Optimal imputation of expression data should improve accuracy of differential expression (DE) analysis. It is a standard practice to benchmark DE calls made on scRNA-Seq data against calls made on their matching bulk counterparts [12]. To this end we used a dataset of myoblasts, for which matching bulk RNA-Seq data were also available [35]. For simplicity this dataset has been referred to as the Trapnell dataset. DE and non-DE genes were identified using edgeR [42] package in R.

We used the standard Wilcoxon Rank-Sum test for identifying differentially expressed genes from matrices imputed by various methods. Congruence between bulk and single cell based DE calls were summarized using the Area Under the Curve (AUC) values yielded from the Receiver Operating Characteristic (ROC) curves (figure 3.(d)). Among all the methods mcImpute performed best with an AUC of 0.85.

For each method, the AUC value was computed on the identical set of ground truth genes. We had to make an exception only for drImpute as it applies the filter to prune genes in its pipeline. Hence AUC value for drImpute was computed based on a smaller set of ground truth genes.

### Improvement in cell type separability

Downstream analysis becomes much easier if expression similarities between cells of identical type are considerably higher than that of cells coming from different subpopulations. To this end, we define cell-type separability score as follows:

For any two cell groups, we first find the median of Spearman correlation values computed for each possible pair of cells within their respective groups. We call the average of the median correlation values the intra-cell type scatter. On the other hand, inter-cell type scatter is defined as the median of Spearman correlation values computed for pairs such that in each pair, cells belong to two different groups. The difference between the intra-cell scatter and inter-cell type scatter is termed as the cell-type separability (CTS) score. We computed CTS scores for two sample cell-type pairs from each dataset. In more than 80 % (13 out of 16) of test cases, mcImpute yielded significantly better CS values (figure 3.(e)-(h), **Table S4**).

### Cell visualization

Representing scRNA-seq data visually would involve reducing the gene-expression matrix to a lower dimensional space and then plotting each cell transcriptome in that reduced two or three dimensional space. Two well-known techniques for dimensionality reduction are PCA and t-SNE [9, 22]. It has been shown that t-Distributed Stochastic Neighbor Embedding (t-SNE) is particularly well suited and effective for the visualization of high-dimensional datasets [20]. So, we use t-SNE (figure 4) on Usoskin and Zeisel expression matrices to explore the performance of dimensionality reduction, both without and with imputation. The cells are visualized in 2-dimensional space, coloring each subpopulation by its annotated group, both before and after imputation. To quantify the groupings of cell transcriptomes, we use an unsupervised clustering quality metric, silhouette index. The average silhouette values for each method have been shown in the plot titles (figure 4).

**Fig 4.**
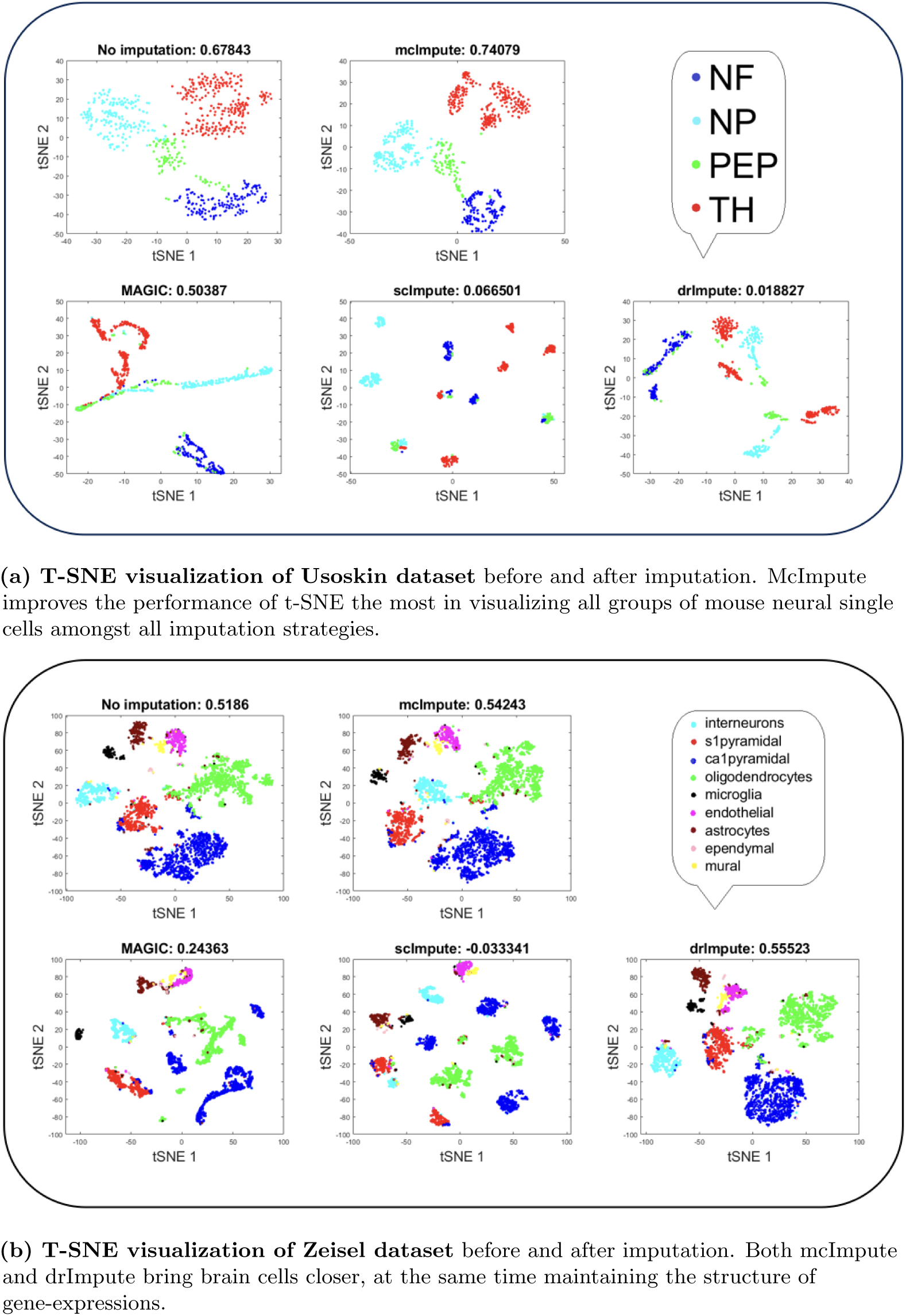
Plots showing [t-SNE visualization: average silhouette values] for (a)Usoskin and (b)Zeisel datasets before and after imputation. McImpute significantly improves the visual distinguishability of scRNA-seq datasets in 2-dimensional space in comparison to other methods.

T-SNE analysis depicts that mcImpute brings all four groups of mouse neural cells from Usoskin data closest to each other in comparison to other methods and performs fairly well, competing with drImpute on Zeisel data too.

### Improvement in distribution of genes

It has been shown that for single-cell gene expression data, in the ideal condition all genes should obey *CV* = *mean*^−1/2^ [44] (CV: coefficient of variation), following a Poisson distribution as depicted by the green diagonal line **(figure 5)**. This is because individual transcripts are sampled from a pool of available transcripts for CEL-Seq. This accounts for technical noise component which obeys Poissonian statistics [45], and thus the CV is inversely proportional to the square root of the mean.

**Fig 5.**
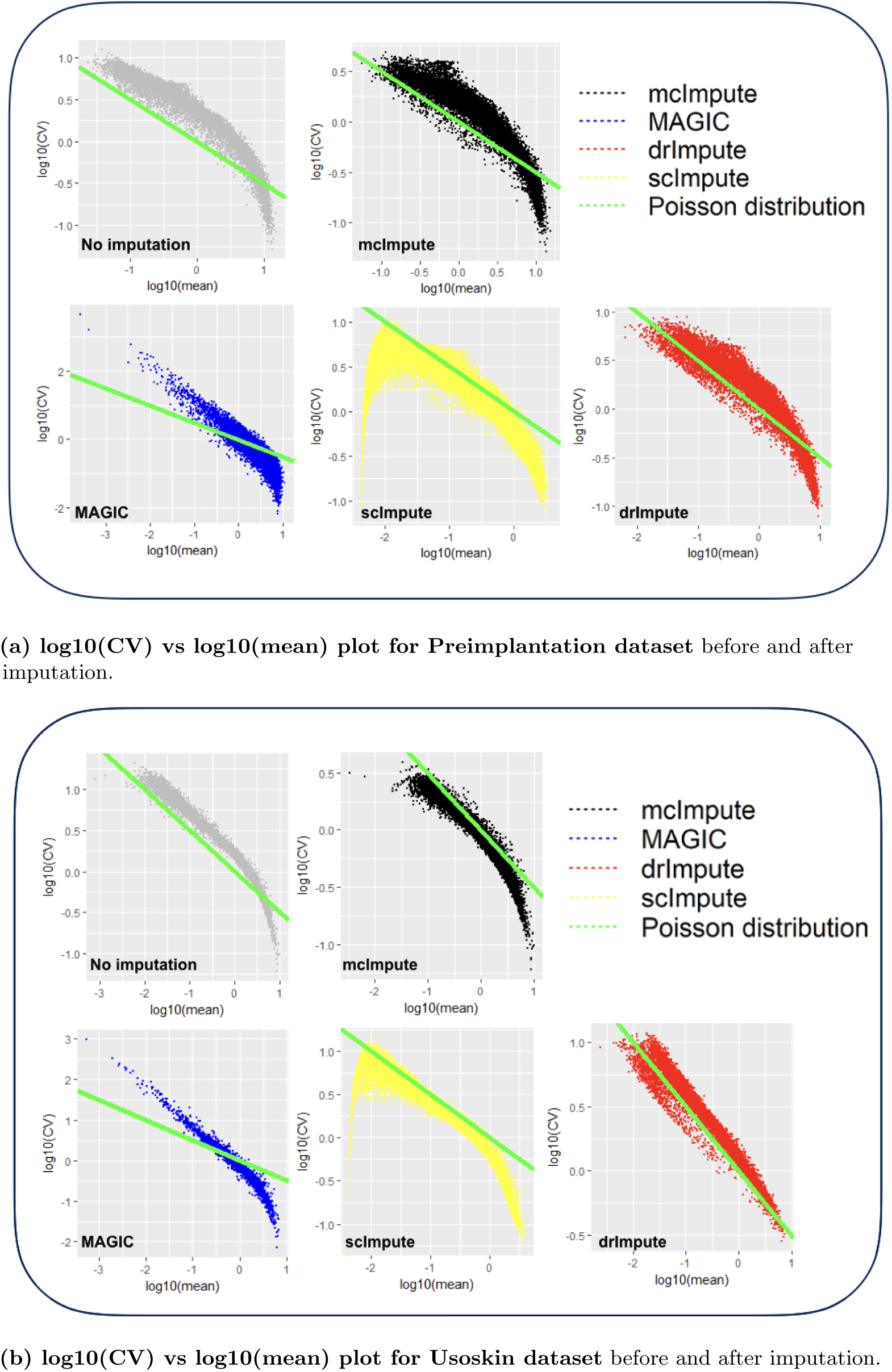
Plots showing log10(CV) vs log10(mean) relationship between genes for (a) Preimplantation and (b) Usoskin datasets before and after imputation. McImpute and drImpute improves the gene distribution, restoring the relationship between CV and mean to be more consistent with Poisson expression distribution, as expected.

We model CV as a function of mean expression for all genes to analyze how various imputation methods affect the relationship between them. The results **(figure 5)** show that both mcImpute and drImpute succeeed to restore the relationship between CV and mean to a great extent (improving the dependency of the CV on the mean expression level to be more consistent with Poissonian sampling noise), while others do not.

## Conclusion

As an inevitable consequence of steep decline in single cell library depth, dropout rates in scRNA-seq data have skyroketed. This works as a confounding factor [8], thereby hindering cell clustering and further downstream analyses. The proposed mcImpute algorithm shows remarkable performance on a number of measures including clustering accuracy, cell type separability, differential gene prediction, cell visualization, gene distribution, etc. McImpute can serve as a very crucial component in single-cell RNA seq pipeline.

Currently imputation and clustering are together a piecemeal two step process - imputation followed by clustering. In the future, we would like to incorporate both clustering and imputation as a joint optimization problem.

## Materials and methods

### Dataset description

We used five scRNA-seq datasets from four different studies for performing various experiments.

- **Jurkat-293T:** This dataset contains expression profiles of Jurkat and 293T cells, mixed *in vitro* at equal proportions (50:50). All ~ 3,300 cells of this data are annotated based on the expressions of cell-type specific markers [41]. Cells expressing CD3D are assigned Jurkat, while those expressing XIST are assigned 293T. This dataset is also available at 10x Genomics website.
- **Preimplantation:** This is an scRNA-seq data of mouse preimplantation embryos. It contains expression profiles of ~ 300 cells from zygote, early 2-cell stage, middle 2-cell stage, late 2-cell stage, 4-cell stage, 8-cell stage, 16-cell stage, early blastocyst, middle blastocyst and late blastocyst stages. The first generation of mouse strain crosses were used for studying monoallelic expression. We downloaded the count data from Gene Expression Omnibus (GSE45719) [40].
- **Zeisel:** Quantitative single-cell RNAseq has been used to classify cells in the mouse somatosensory cortex (S1) and hippocampal CA1 region based on 3005 single cell transcriptomes [43]. Individual RNA molecules were counted using unique molecular identifiers (UMIs) and confirmed by single-molecule RNA fluorescence in situ hybridization (FISH). A divisive biclustering method based on sorting points into neighborhoods (SPIN) was used to discover molecularly distinct, 9 major classes of cells. Raw data is available under the accession number GSE60361.
- **Usoskin:** This data of mouse neurons [37] was obtained by performing RNA-Seq on 799 dissociated single cells dissected from the mouse lumbar dorsal root ganglion (DRG) distributed over a total of nine 96-well plates. The cell labels (clusters of mouse lumbar DRG-NF, NP, TH, PEP populations) were computationally derived and assigned by performing PCA classification on single mouse neurons. RPM normalized counts are available under the accession number GSE59739.
- **Trapnell**: This is an scRNA-seq data of primary human myoblasts [35]. Differentiating myoblasts were cultured and cells were dissociated and individually captured at 24-hour intervals. 50–100 cells at each of four time points were captured following serum switch using the Fluidigm C1 microfluidic system. This data is available at Gene Expression Omnibus under the accession number GSE52529.

### Data preprocessing

Steps involved in preprocessing of raw scRNA-seq data are enumerated below.

- **Data filtering:** It is ensured that data has no bad cells and if a gene was detected with ≥ 3 reads in at least 3 cells we considered it expressed. We ignored the remaining genes.
- **Library-size Normalization:** Expression matrices were normalized by first dividing each read count by the total counts in each cell, and then by multiplying with the median of the total read counts across cells.
- **Log Normalization:** A copy of the matrices were log_2_ transformed following addition of 1 as pseudocount.
- **Imputation:** Further, log transformed expression matrix was used as input to mcImpute. The algorithm returns imputed log transformed matrix, normalized matrix (after anti-log operation on imputed log-transformed expressions) and the count matrix after imputation.

### Is a gene expression matrix low-rank?

Expression levels of genes at a particular instance is orchestrated by a complex regulatory network. This interdependency is best reflected by pairwise correlations between genes. It has previously been argued that a small number of interdependent biophysical functions trigger the functioning of transcription factors, which in turns influence the expression levels of genes, resulting in a highly correlated data matrix [10]. On the other hand, cells coming from same tissue source also lie on differential grades of variability of a limited number phenotypic characteristics. Therefore, it is just to assume that the gene expression values lie on a low-dimensional linear subspace and the data matrix thus formed may well be thought as a low-rank matrix.

### Low-rank matrix completion: Definition

Our problem is to complete a partially observed gene expression matrix *X* where columns represent genes and rows, individual cells. The complete matrix is constituted by the known and the yet unknown values. We can assume that the single cell data that we have acquired, *Y* is a sampled version of the complete expression matrix *X*. Mathematically, this is expressed as,

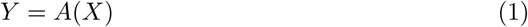

Here *A* is the sub-sampling operator. It is a binary mask that has 0’s where the counts of complete expression data *X* has not been observed and 1’s where they have been. Our problem is to recover *X*, given the observations *Y*, and the sub-sampling mask *A*. It is known that *X* is of low-rank.

It should be noted that matrix completion is a well studied framework. In this work, we propose two algorithms for efficient imputation of scRNA-seq expression data-Matrix factorization and Nuclear norm minimization.

### Matrix factorization

Matrix factorization is the most straightforward way to address the low-rank matrix completion problem; it has previously been used for finding lower dimensional decompositions of matrices [17]. Say *X* is of dimensions *m* × *n*, but is known to have a rank *r* (<*m, n*). In that case, one can express *X_m×n_* as a product of two matrices *U_m×r_* and *V_r×n_*. Therefore the complete problem (1) can be formulated as,

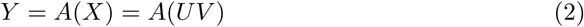

Estimating *U* and *V* from (2) tantamount to recovering *X*. The two matrices *U* and *V* can be solved by minimizing the *Frobenius* norm of the following cost function.

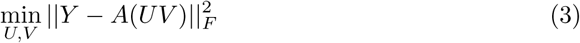

Since this is a bi-linear problem, one cannot guarantee global convergence. However it usually works in practice. It has been used for solving recommender systems problems [13], where (3) was solved using stochastic gradient descent (SGD). SGD is not an efficient techniques and requires tuning of several parameters. In this work, we will solve (3) in a more elegant fashion using majorization minimization [31]. The basic MM approach and its geometrical interpretation has been diagrammatically represented (**figure S1**). It depicts the solution path for a simple scalar problem but essentially captures the MM idea.

For our given problem, 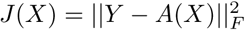 the majorization step basically decouples the problem (from A), so that we can solve the optimization problem by solving

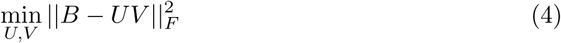

where *B* = *X_k_* + *A^T^*(*Y* − *A*(*X_k_*)) at each iteration k.

This (4) is solved by alternating least squares, i.e. while updating *U*, *V* is assumed to be constant and while updating *V*, *U* is assumed to be constant.

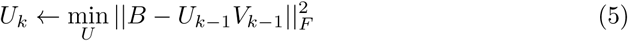

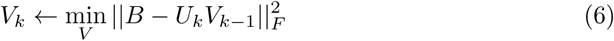

Since the log-transformed input (with pseudo count added) expressions would never be negative, we have imposed non-negativity constraint on the recovered matrix X, so that it does not contain any negative values.

The matrix factorization algorithm has been summarized in algorithm 1. The initialization of factor V is done by keeping *r* right singular vectors of X in V, where *r* is the approximate rank of the expression matrix to be recovered.

#### Algorithm 1 Matrix completion using matrix factorization

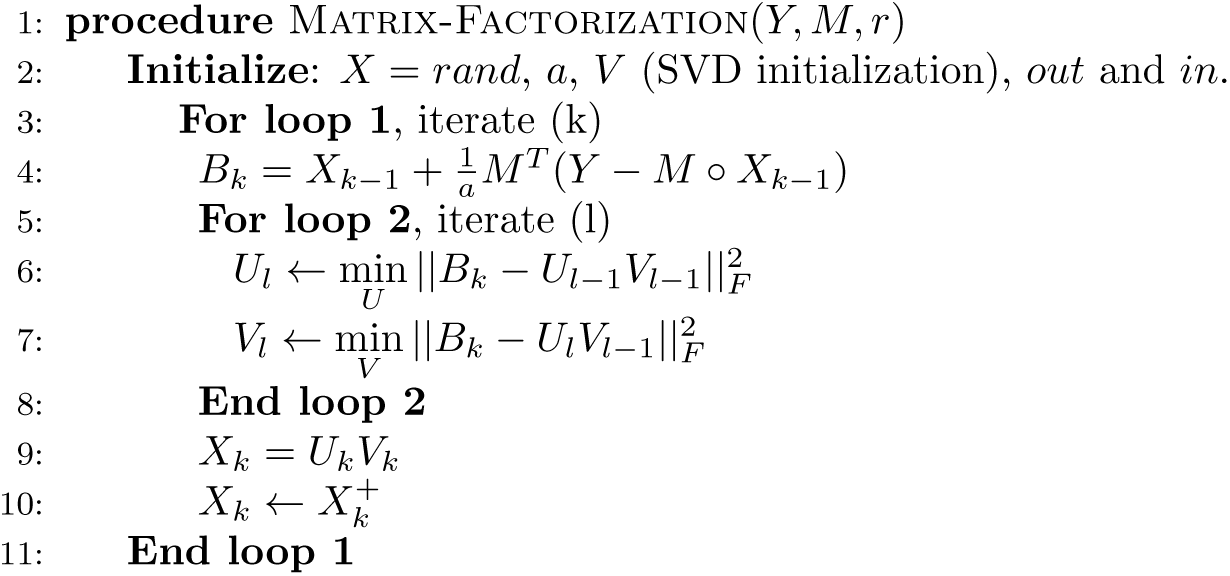

### Nuclear norm minimization

The problem depicted in (3) is non-convex. Hence, there is no guarantee for global convergence. Also one needs to know the approximate rank of the matrix *X* in order to solve it, which is unknown in this case. To combat this issues, researchers in applied mathematics and signal processing proposed an alternative solution. They would directly solve the original problem (1) with a constraint that the solution is of low-rank. This is mathematically expressed as,

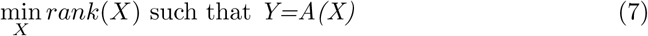

However this turns out to be NP hard problem with doubly exponential complexity. Therefore, studies in matrix completion [6, 7] proposed relaxing the NP hard rank minimization problem to its closest convex surrogate: nuclear norm minimization.

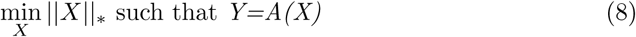

Here the nuclear norm is defined as the sum of singular values of data matrix *X*. It is the l1 norm of the vector of singular values of X and is the tightest convex relaxation of the rank of matrix, and therefore its ideal replacement.

This is a semi-definite programming (SDP) problem. Usually its relaxed version (Quadratic Program) is solved [5].

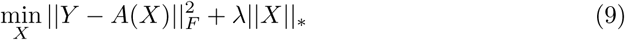

The problem (9) does not have a closed form solution and needs to be solved iteratively.

To solve (9), we invoke MM once more. Here 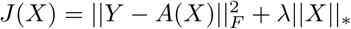, we can express (9) in the following fashion in every iteration *k*

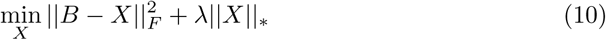

where *B* = *X_k_* + *A^T^*(*Y* − *A*(*X_k_*)).

Using the inequality ||*Z*_1_ − *Z*_2_||*_F_* ≥ ||*s*_1_ − *s*_2_||_2_, where *s*_1_ and *s*_2_ are singular values of the matrices *Z*_1_ and *Z*_2_ respective, we can solve the following instead of solving the minimization problem (10).

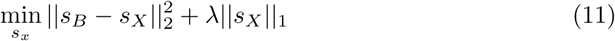

Here *s_B_* and *s_X_* are the singular values of *B* and *X* respectively. It has been shown that problem (10) is minimized by soft thresholding the singular values with threshold *λ*/2. The optimal update is given by

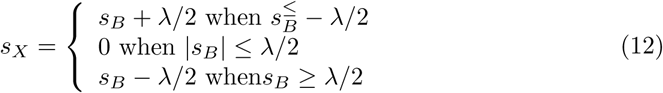

or more compactly by

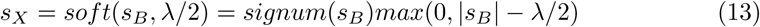

#### Algorithm 2 Matrix completion via iterated soft thresholding

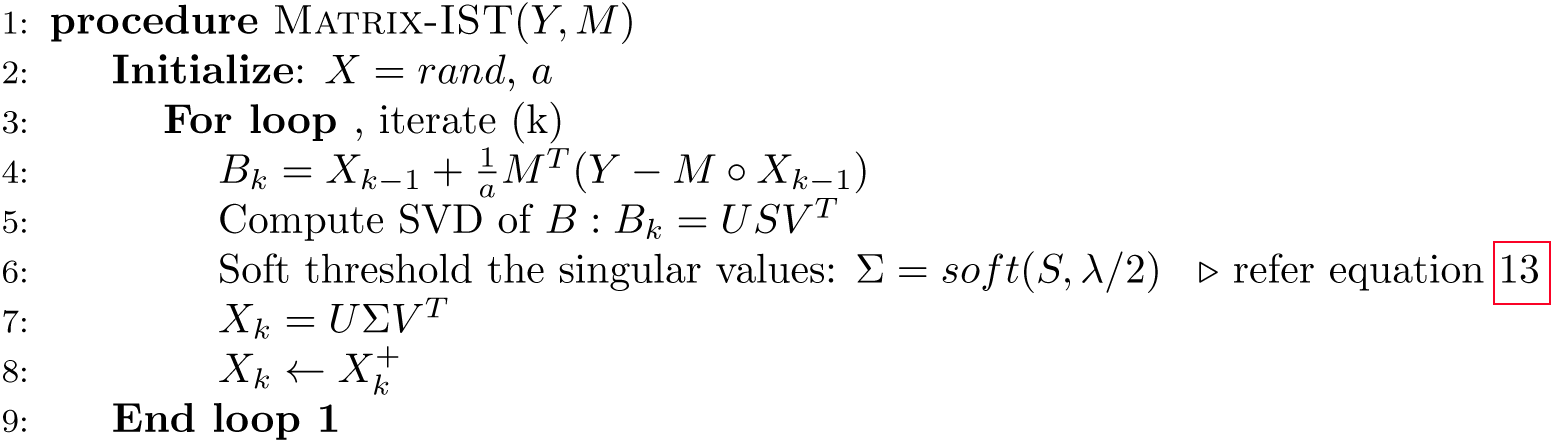

We found that the algorithm is robust to values of λ as long as as it is reasonably small (< 0.01).

Here too, we have imposed non-negativity constraint on *X* since expressions cannot be smaller than zero.

## Supporting information

S1 Fig. Schemematic diagram of Majorization-Minimization.

**S1 Table. Separation of “true zeros” from dropouts.** Fraction of zeros (values between 0 and 0.5) in single cell expression matrix against the median bulk expression. The genes are divided into 10 bins based on median bulk genes expression (first bin has only 0 expression gene

**S2 Table. Matrix recovery error.** Comparison of NMSE, RMSE and MAE between recovered matrices and unmasked Usoskin data using Nuclear Norm Minimization (NNM) and Matrix Factorization (MF) algorithms at hidden/masked positions

**S3 Table. Average Adjusted Rand Index values.** Average Adjusted Rand Index values (on 100 runs of PCA followed by k-means) measuring the correspondence between the k-means predicted clusters and the prior annotations.

**S4 Table. CTS scores:.** Cell type separability (CTS) for any 2 randomly chosen cell groups from each dataset.

## Software

The source code of mcImpute is shared at https://github.com/aanchalMongia/McImpute_scRNAseq

## Acknowledgments

This work was supported by a grant by DST/INT/CANADA/IC-IMPACT/P-8/2016 awarded to AM and INSPIRE Faculty Award grant DST/INSPIRE/04/2015/003068 given to DS by DST, Govt. of India.

